# Increased Binding of [^18^F]Nifene, a PET Imaging Probe for α4β2* Nicotinic Acetylcholinergic Receptors in Hippocampus-Subiculum of Postmortem Human Parkinson’s Disease Brain

**DOI:** 10.64898/2026.04.07.716971

**Authors:** Jogeshwar Mukherjee, Fariha Karim, Allyson Ngo, Christopher Liang, Geidy E. Serrano, Thomas G. Beach

**Affiliations:** Preclinical Imaging, Department of Radiological Sciences, University of California-Irvine, Irvine, CA 92697, USA; Banner Sun Health Research Institute, Sun City, AZ 85351, USA

**Keywords:** α−Synucleinopathy, Lewy Bodies, PET, Nifene, Neurodegeneration

## Abstract

Non-motor symptoms in Parkinson’s disease (PD) may be influenced by the α4β2* subtype of nicotinic acetylcholine receptors (nAChR) present in the hippocampus and subiculum. To continue efforts in PET diagnostics for PD, autoradiographic [^18^F]nifene binding to α4β2* nAChR was quantitively assessed in the hippocampus-subiculum (HP-SUB) of PD (n = 27; 14 males, 13 females) and cognitively normal (CN) (n = 32; 16 males, 16 females) cases. Anti-ubiquitin for Lewy body and anti-α-synuclein immunostaining on adjacent slices were analyzed in QuPath and [^18^F]nifene binding was quantified in OptiQuant. Subiculum had greater [^18^F]nifene binding (51% to 85%) compared to HP in all subjects. Significantly higher [^18^F]nifene binding (>250%) was seen in PD SUB and PD HP compared to CN in both males and females. The grey matter (GM) to white matter (WM) ratio in PD=3.53 while CN=1.33, a >150% increase in PD. Binding of [^18^F]nifene to GM and WM individually was >250% greater in PD compared to CN. Male CN exhibited an increase while and male PD exhibited a significant decrease in [^18^F]nifene binding with aging, while females did not exhibit significant differences. In summary, α4β2* nAChR measured by [^18^F]nifene is significantly upregulated in the PD HP and SUB. This increased [^18^F]nifene binding may be of diagnostic value using PET imaging.

## 1 Introduction

Non-motor symptoms in Parkinson’s disease (PD) including cognitive deficits, psychosis, and depression are associated with the somato-cognitive action network (SCAN) (Robbins & Cools 2014; Lenka et al. 2016; <u>Hussein et al. 2021</u>; Ren et al., 2026). The prominence of non-motor symptoms can arise early in PD, commonly preceding motor symptoms (<u>Hussein et al. 2021</u>; Silva et al. 2023). The severity of these symptoms can be determined by pathological abnormalities in PD. The accumulation of α-synuclein is sufficient to initiate a neurodegenerative cascade resulting in the loss of dopaminergic neurons, especially the substantia nigra, and motor impairment (Luk et al. 2012). The loss of dopaminergic neurons throughout the brain is accompanied by the rise of the protein deposits Lewy bodies. Intraneuronal Lewy bodies contain misfolded fibrillar α-synuclein aggregates that damage neurons and mitochondria in pathways that contribute to cell death, neuroinflammation and neurodegeneration (Bandopadhyay & Belleroche 2010; Henrich et al. 2023; Mukherjee et al., 2022). Membrane permeability is increased by α-synuclein protofibrils, dysregulating calcium signaling required for neurotransmission and autophagy (Angelova et al. 2016; Post et al. 2018). Due to the numerous dysfunctional proteins in Lewy bodies, they are ubiquitinated alongside misfolded α-synuclein but the continuous accumulation of ubiquitinated proteins overwhelms the proteolytic degradation system (Tofaris et al. 2003).

All cells in the dopaminergic system have nicotinic acetylcholine receptors (nAChR) that promote cognition via neuroprotection and synaptic plasticity (Iarkov et al. 2021). Both the α4β2* and α6β2 nAChR subtypes mediate tonic dopamine release, therefore maintaining striatal dopaminergic tone (Perez et al. 2010). The α4β2* and α7 nAChR subtypes have been suggested to play a role in dopamine neuron survival, mitochondrial dysfunction, cognitive impairment, and neuroprotection (Guan 2024; Lin et al. 2025). Amongst the abundant receptors in the central nervous system (CNS), the α4β2* subtype is involved in the maintenance of cognitive function in brain regions including the amygdala, hippocampus (HP), striatum, and cerebellum (Picciotto et al. 2000; Nees 2015; Bieszczcad et al., 2012). The hippocampus is majorly responsible for learning and memory formation, so hippocampal neurogenesis is especially vulnerable to neurodegeneration and cognitive decline (Terreros-Roncal et al. 2021). The dense population of α4β2 nAChR* in the hippocampus modulates cell activity and plasticity while also enhancing learning and contextual fear conditioning in the presence of intrahippocampal nicotine (Davis et al. 2007; Laikowski et al. 2019). Input modulation by multiple nAChR subtypes in the subiculum (SUB), an interface between the hippocampus and the brain reward circuitry, includes indirect activation of dopamine neuronal activity which is crucial for locomotion and learning (Graham et al. 2003; Cooper et al. 2006). In addition to changes in nAChRs, both α-synuclein aggregates and Lewy bodies undergo changes in PD that can affect functions in the HP-SUB (Figure 1).

**Figure 1.**
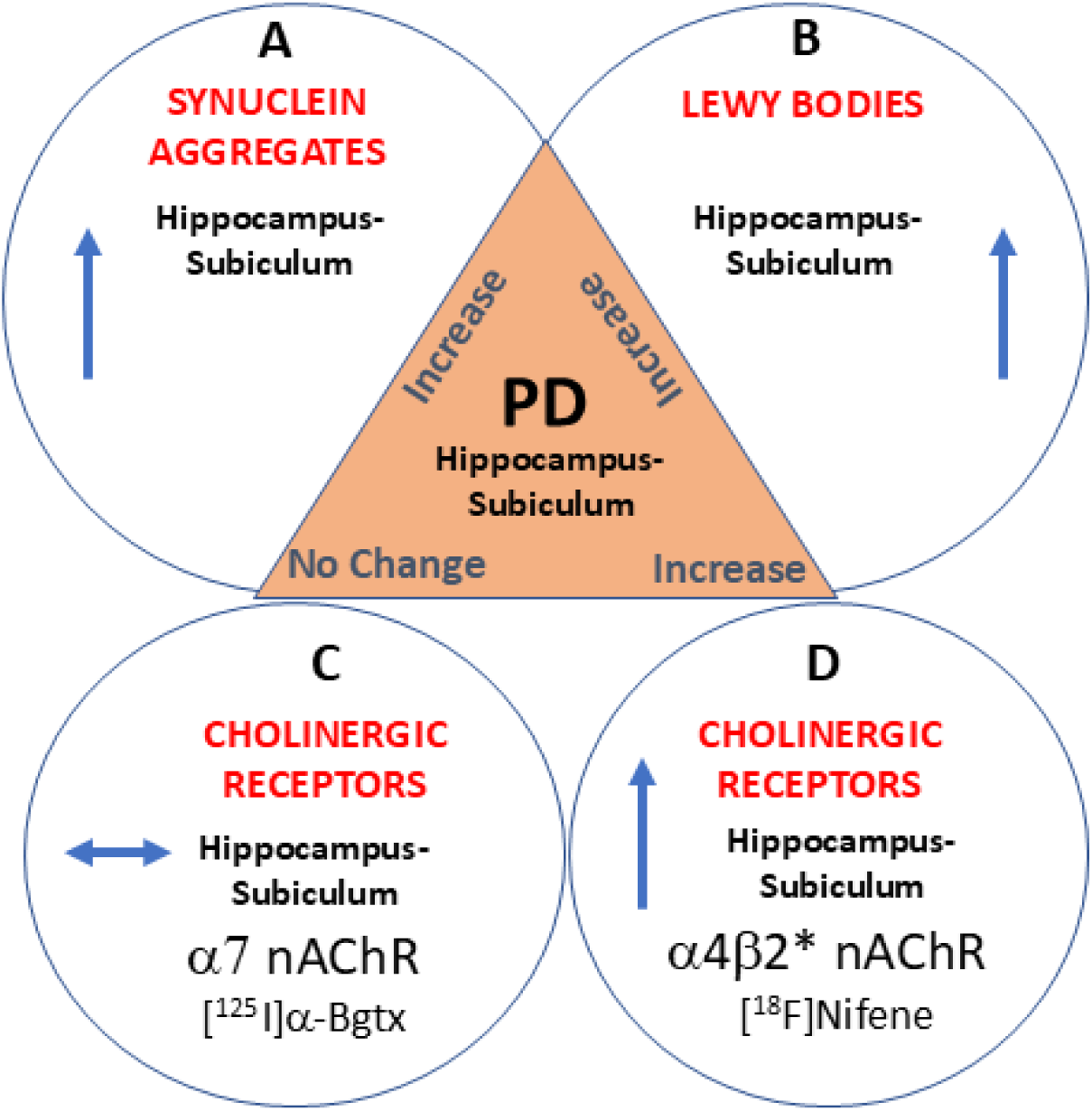
Schematic showing potential changes in nAChRs in PD. (**A**). α-Synuclein aggregates are increased and may affect nAChRs in PD; (**B**). Increased Lewy bodies accumulation may cause nAChRs alterations in PD; (**C**). [^125^I]α-Bgtx binding to α7 nAChRs in this cohort of postmortem PD brains was not affected in HP-SUB; (**D**). [^18^F]Nifene binding in HP-SUB in this cohort of postmortem PD brains was elevated as reported in this study.

The relevance of cholinergic pathways in PD symptoms has been suggested in numerous positron emission tomography (PET) studies. The non-invasive nature of PET imaging α4β2 nAChR allows for the sensitive detection of abnormalities in neurodegenerative diseases including PD (Tiepolt et al. 2022). An early tracer for α4β2* nAChR, 5-[^125^I]-A-85380, found an in vitro alignment between the loss of nigrostriatal dopaminergic markers and striatal α4β2* in PD (Pimlott et al. 2004). The single-photon emission computed tomography (SPECT) equivalent, 5-[^123^I]-A-85380, agreed with high uptake in the thalamus while also correlating cerebral α4β2* nAChR density with PD associated cognitive dysfunctions (Lorenz et al. 2014). Reduction of 2-[^18^F]FA-85380 binding to α4β2* nAChR associated with the severity of mild cognitive or depressive symptoms in PD subjects within the subcortical and cortical regions, substantia nigra, and striatum (Meyer et al. 2009; Kas et al. 2009). The PET tracer 2-[^18^F]FA-85380 was commonly used for in vivo examinations of α4β2* nAChR but is limited by slow kinetics and may differ in the degree of reductions within particular brain regions (Tiepolt et al. 2022). [^18^F]XTRA PET and 5-[^123^I]-A-85380 SPECT binding to α4β2* nAChR was higher in PD compared to control, contradicting other PET studies that confirm a decrease in PD subjects (Mills et al. 2025; Isaias et al. 2014). Discrepancies may be due to differences among the PET tracers, imaging characteristics and affinity dependent on the brain region and stage of PD (Mills et al. 2025).

[^18^F]Nifene is a moderate affinity analog of 2-[^18^F]FA-85380 with more rapid kinetic properties than aforementioned PET agents (Pichika et al. 2006; Hillmer et al. 2012). Human PET studies with [^18^F]nifene have been successful in quantitative assessments of α4β2* nAChR and brain regional connectivity, indicating suitability for CNS disorders (Lao et al. 2017; Mukherjee et al. 2018). A quantitative autoradiographic evaluation of [^18^F]nifene on Alzheimer’s disease (AD) hippocampal brain sections indicated a significant reduction of α4β2* nAChR compared to cognitively normal (CN) (<u>Karim et al. 2025</u>). The success of this evaluation and previous implications of the cholinergic pathway in PD prompted a similar assessment of [^18^F]nifene binding to α4β2* nAChR in PD. The purpose of this study was to assess in vitro [^18^F]nifene binding to α4β2* nAChR in human PD postmortem hippocampus (HP)-subiculum (SUB) brain slices. Findings may indicate unique features of the cholinergic effects on PD when compared to CN, justifying the use of [^18^F]nifene in PD including PET studies.

## 2 Methods

### 2.1 Postmortem Human Brains

Human postmortem brain tissue samples were obtained from Banner Sun Health Research Institute (BHRI), Sun City, AZ, brain tissue repository for in vitro experiments. Detailed characteristics were given with each sample (Table 1). Brain tissue samples from PD (n=27, 14 male and 13 female) and CN (n=32, 16 male and 16 female) were selected for the presence and absence of end-stage pathology. AD cases were previously reported (<u>Karim et al. 2025</u>). All brain slices of each case contained HP and SUB regions. Brain sections were stored at −80°C. All postmortem human brain studies were approved by the Institutional Biosafety Committee of University of California, Irvine.

**Table 1.**
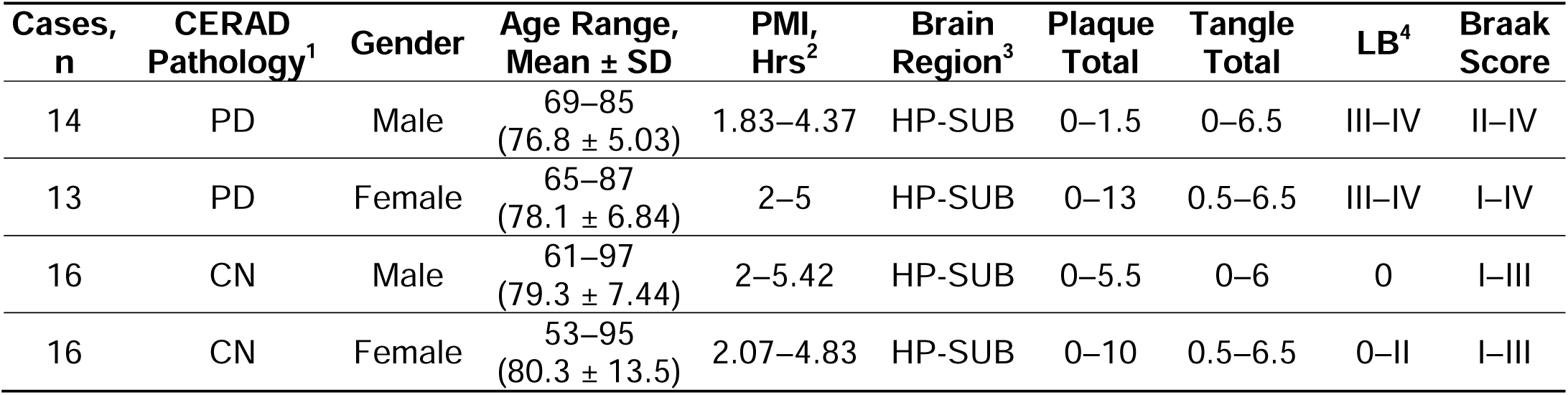
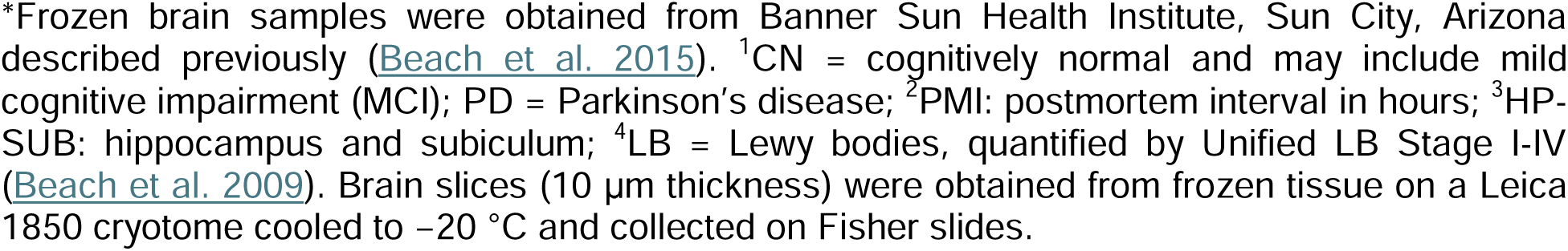
Patient samples and data*.

| Cases, n | CERAD Pathology<sup>1</sup> | Gender | Age Range, Mean ± SD | PMI, Hrs<sup>2</sup> | Brain Region<sup>3</sup> | Plaque Total | Tangle Total | LB<sup>4</sup> | Braak Score |
| --- | --- | --- | --- | --- | --- | --- | --- | --- | --- |
| 14 | PD | Male | 69–85<br>(76.8 ± 5.03) | 1.83–4.37 | HP-SUB | 0–1.5 | 0–6.5 | III–IV | II–IV |
| 13 | PD | Female | 65–87<br>(78.1 ± 6.84) | 2–5 | HP-SUB | 0–13 | 0.5–6.5 | III–IV | I–IV |
| 16 | CN | Male | 61–97<br>(79.3 ± 7.44) | 2–5.42 | HP-SUB | 0–5.5 | 0–6 | 0 | I–III |
| 16 | CN | Female | 53–95<br>(80.3 ± 13.5) | 2.07–4.83 | HP-SUB | 0–10 | 0.5–6.5 | 0–II | I–III |

\*Frozen brain samples were obtained from Banner Sun Health Institute, Sun City, Arizona described previously (Beach et al. 2015). <sup>1</sup>CN = cognitively normal and may include mild cognitive impairment (MCI); PD = Parkinson's disease; <sup>2</sup>PMI: postmortem interval in hours; <sup>3</sup>HP-SUB: hippocampus and subiculum; <sup>4</sup>LB = Lewy bodies, quantified by Unified LB Stage I-IV (Beach et al. 2009). Brain slices (10 µm thickness) were obtained from frozen tissue on a Leica 1850 cryotome cooled to –20 °C and collected on Fisher slides.

### 2.2 Radiopharmaceuticals

Fluorine-18 fluoride in oxygen-18 enriched water was purchased from PETNET, Inc. Fluorine-18 radioactivity was counted in a Capintec CRC-15R dose calibrator while low level counting was carried out in a Capintec Caprac-R well-counter. The radiosynthesis of [^18^F]nifene was performed using nucleophilic displacement of the nitro group in *N*-BOC-nitronifene precursor (Pichika et al., 2006) by [^18^F]fluoride in an automated synthesizer followed by deprotection using previously described procedures (Pichika et al., 2006, Campoy et al., 2021, Bhuiyan et al., 2024). All solvents used were provided by Fisher Scientific. Reactions were monitored using analytical thin-layer chromatography (TLC) (Baker-flex, Phillipsburg, NJ, USA). RadioTLC were scanned on an AR-2000 imaging scanner (Eckart & Ziegler, Berlin, Germany). Radiochemical purity of [^18^F]nifene was >98% and chemical purity was found to be >95% with a measured molar activity >70 GBq/μmol (>2 Ci/μmol) at the end of synthesis.

### 2.3 [^18^F]Nifene Autoradiography

Brain slices of PD and CN cases on glass slides were placed into glass chambers and preincubated in Tris buffer (50 mmol/L Tris containing 120 mmol/L NaCl, 5 mmol/L KCl, 2.5 mmol/L CaCl_2_, 1 mmol/L MgCl_2_, pH 7.4) for 15 min. After discarding the preincubation buffer, [^18^F]nifene in Tris buffer was added to each chamber for incubation at 25°C for 1 hour. The slides underwent a series of washing with cold buffer for 3 minutes twice and cold deionized water for one minute. The slides were air dried and placed into a film cassette with a phosphor screen film. The next day, the films were taken out of the cassettes and read using the Cyclone Phosphor Imaging System (Packard Instruments Co) to produce autoradiographic images. The images were opened on the OptiQuant acquisition and analysis program (Packard Instruments Co) to draw regions of interest and measure [^18^F]nifene binding measured in digital light units/mm^2^ (DLU/mm^2^).

### 2.4 Immunohistochemistry

Adjacent slices of each case were immunnostained with anti-ubiquitin and anti-α-synuclein of all brain slices was carried out by University of California-Irvine, Pathology services using Ventana BenchMark Ultra protocols. Neighboring slices were immunostained for Ubiquitin (Cell Marque catalog no. 318A-18, Rocklin, CA, USA) and α-synuclein (EMD Millipore Corporation, lot No. 2985418, Burlington, MA, USA). All immunohistochemistry (IHC) slides were scanned using the Ventana Roche instrumentation and analyzed using QuPath (version QuPath-0.4.2).

### 2.5 Statistical Analysis

GraphPad Prism 10 and Microsoft Excel 16 were used to assess [^18^F]nifene binding quantified by DLU/mm^2^ values from OptiQuant. Statistical power was determined with unpaired two-tailed parametric Student’s t-test, and p values of <0.05 indicated statistical significance. Error bars signify mean□±□standard deviation. The Shapiro–Wilk test confirmed normality of distribution among all groups.

## 3 Results

### 3.1. [^18^F]Nifene in Subiculum and Hippocampus in PD Cases

Figure 2 consists of images of female PD 05-17 as a representative case. The location of the SUB and HP were identified in all female PD cases (Figure 2A). An adjacent brain slice was used to positively immunostain for anti-ubiquitin and anti-α-synuclein to confirm the presence of Lewy bodies and α-synuclein in all female PD (Figure 2C,D). There were greater quantities of Lewy bodies and α-synuclein in the SUB than the HP, aligning with the distribution of [^18^F]nifene binding (Figure 2B). Throughout all female PD cases, the SUB was found to have noticeably greater [^18^F]nifene compared to the HP (Figure 2E).

**Figure 2.**
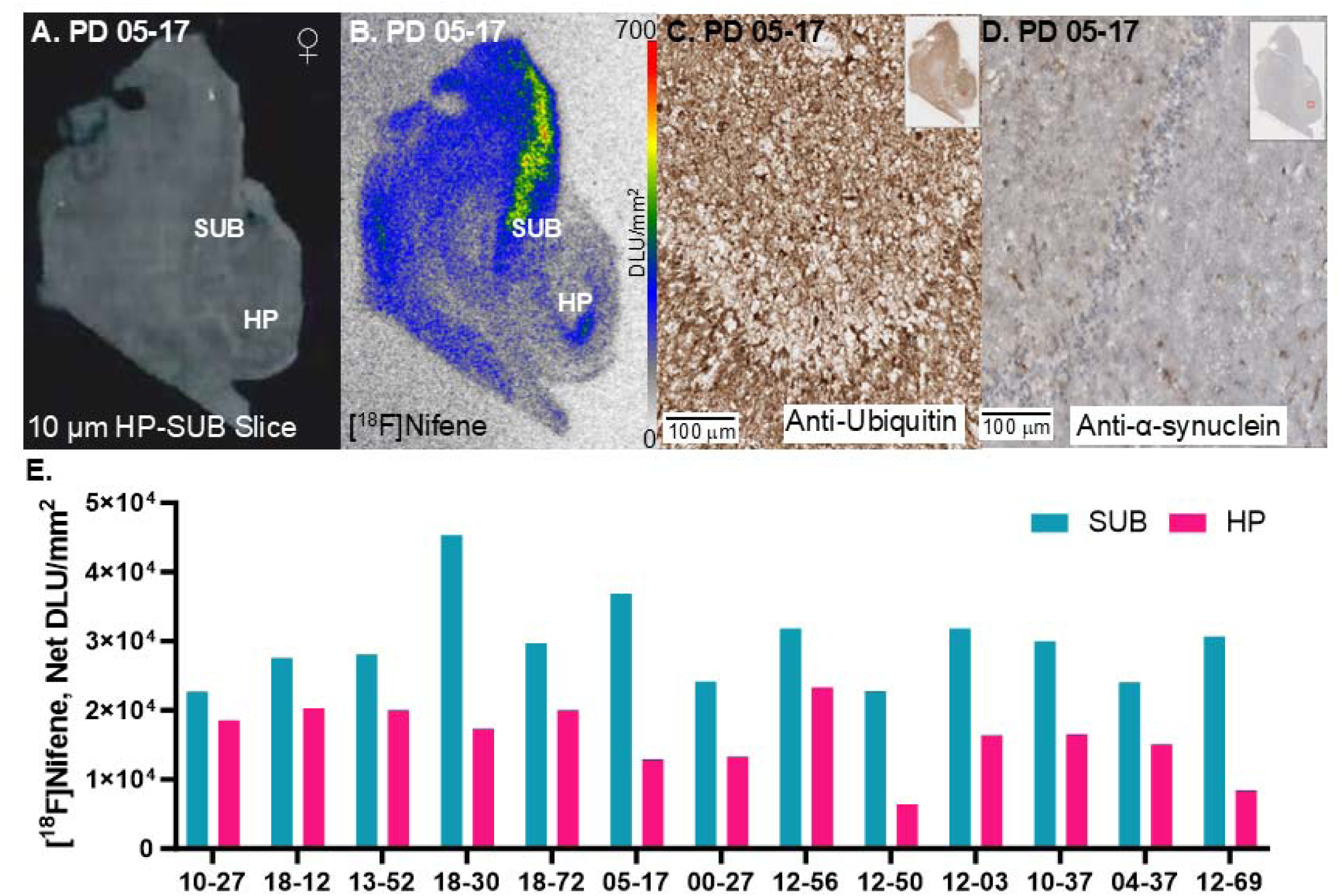
[^18^F]nifene binding to α4β2 nAChR in representative female PD. (A) Scan of postmortem human HP-SUB brain slice (10 μm) mapping location of subiculum (SUB) and hippocampus (HP). (B) Autoradiographic image of PD 05-17 with [^18^F]nifene binding; autoradiography scale bar: 0-700 DLU/mm^2^. (C) IHC image of anti-ubiquitin on PD 05-17; 100 μm magnification. (D) IHC image of anti-α-synuclein on PD 05-17; 100 μm magnification. (E) Quantification of [^18^F]nifene within SUB and HP regions of all PD female.

Figure 3 shows images of male PD 16-65 as a representative case. The location of the SUB and HP were identified in all male PD cases (Figure 3A). An adjacent brain slice was positively immunostain for anti-ubiquitin and anti-α-synuclein to confirm the presence of Lewy bodies and α-synuclein in all male PD (Figure 3C,D). [^18^F]nifene binding to the SUB was substantially greater than the HP in all male PD cases, both visually and quantitatively (Figure 3B,E).

**Figure 3.**
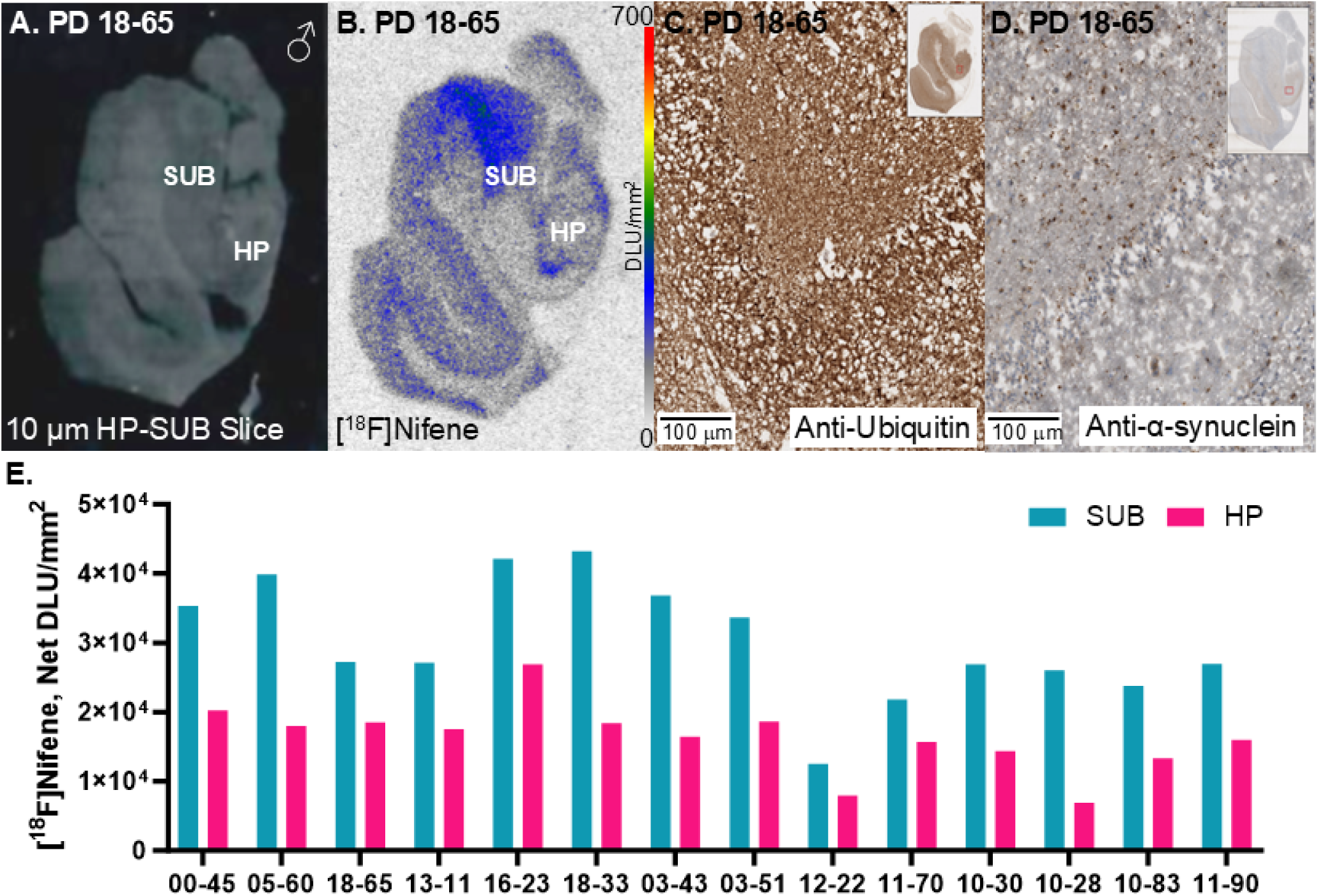
[^18^F]nifene binding to α4β2 nAChR in representative male PD. (A) Scan of postmortem human HP-SUB brain slice (10 μm) mapping location of subiculum (SUB) and hippocampus (HP). (B) Autoradiographic image of PD 18-65 with [^18^F]nifene binding; autoradiography scale bar: 0-700 DLU/mm^2^. (C) IHC image of anti-ubiquitin on PD 18-65; 100 μm magnification. (D) IHC image of anti-α-synuclein on PD 18-65; 100 μm magnification. (E) Quantification of [^18^F]nifene within SUB and HP regions of all PD male.

### 3.2. [^18^F]Nifene in Subiculum and Hippocampus in CN Cases

Figure 4 includes both female and male CN representative cases. Localization of the SUB and HP were identified in all CN cases (Figure 4A). When immunostained for anti-ubiquitin and anti-α-synuclein, there were lower quantities of Lewy bodies and α-synuclein throughout the adjacent brain slices while mostly concentrating within the SUB region (Figure 4C,D,G,H). In all CN cases, [^18^F]nifene binding was greater in the SUB compared to HP (Figure 4B,F,I).

**Figure 4.**
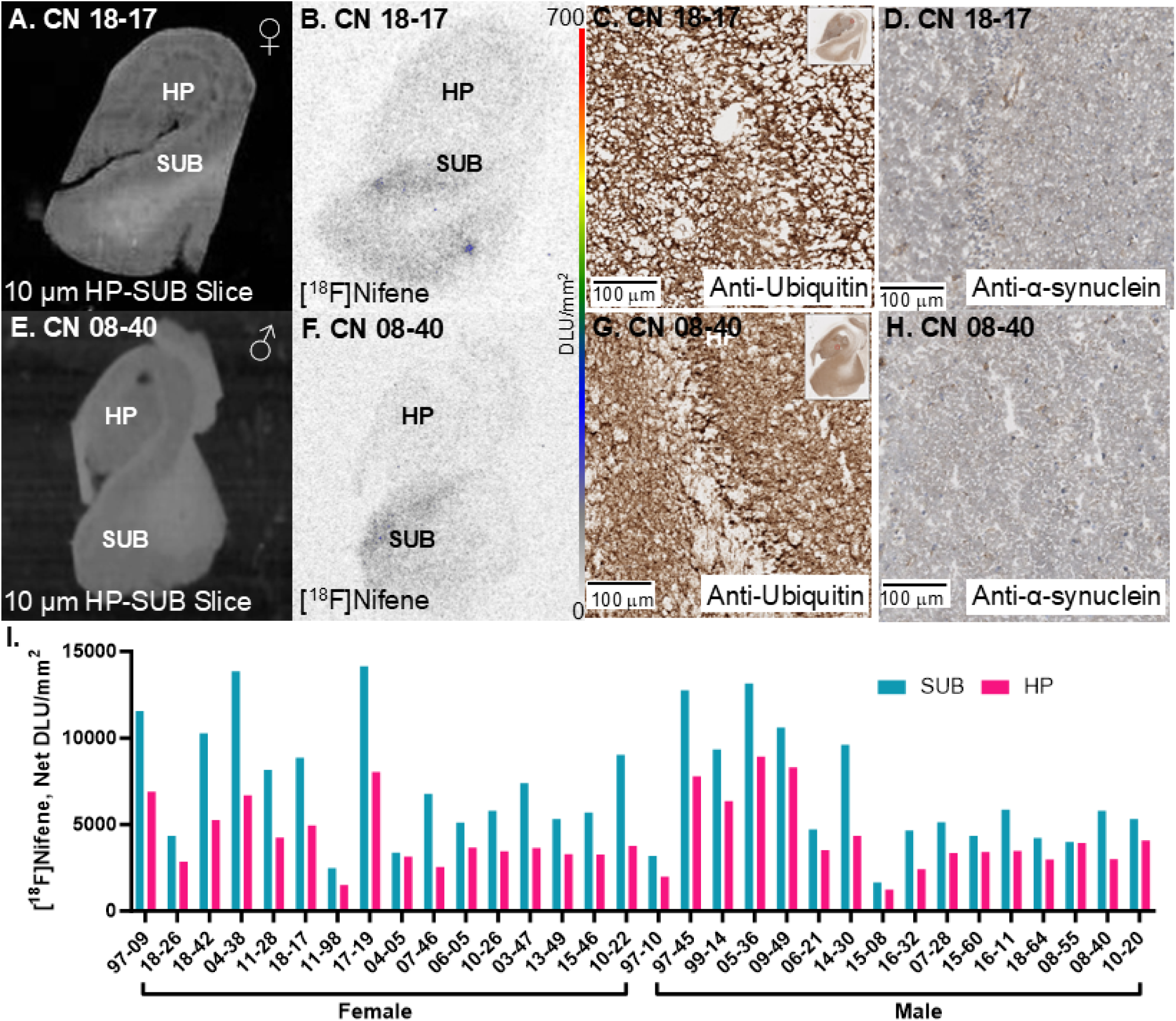
[^18^F]nifene binding to α4β2 nAChR in representative CN. (A) Scan of female CN 18-17 postmortem human HP-SUB brain slice (10 μm) mapping location of subiculum (SUB) and hippocampus (HP). (B) Autoradiographic image of CN 18-17 with [^18^F]nifene binding; autoradiography scale bar: 0-700 DLU/mm^2^. (C) IHC image of anti-ubiquitin on CN 18-17; 50 mm magnification. (D) IHC image of anti-α-synuclein on CN 18-17; 50 mm magnification. (E) Scan of male CN 08-40 postmortem human HP-SUB brain slice (10 μm) mapping location of subiculum (SUB) and hippocampus (HP). (F) Autoradiographic image of CN 08-40 with [^18^F]nifene binding; autoradiography scale bar: 0-700 DLU/mm^2^. (G) IHC image of anti-ubiquitin on CN 08-40; 100 μm magnification. (H) IHC image of anti-α-synuclein on CN 08-40; 100 μm magnification. (I) Quantification of [^18^F]nifene within SUB and HP regions of all CN cases.

### 3.3. [^18^F]Nifene PD and CN Group Comparisons in Subiculum and Hippocampus

Comparisons of [^18^F]nifene binding between SUB and HP were made within PD and CN cases. All comparisons between SUB and HP within PD were significant while CN were not significant (Figure 5A,B,C). In male regional comparisons, [^18^F]nifene binding to SUB was greater than HP in CN by 51% (CN ♂ SUB/HP= 1.51) and in PD by 85% (PD ♂ SUB/HP= 1.85) (Figure 5A). When comparing [^18^F]nifene binding between CN males to PD males, PD SUB was higher than CN SUB by >350% (PD ♂ SUB/ CN ♂ SUB= 4.64) and in the case of HP, PD HP was higher than CN HP by >250% (PD ♂ HP/ CN ♂ HP= 3.78) (Figure 5A). In the case of females, regional comparisons of [^18^F]nifene binding to SUB was also greater than HP in CN by 82% (CN ♀ SUB/HP= 1.82) and in PD by 85% (PD ♀ SUB/HP= 1.85) (Figure 5B). When comparing [^18^F]nifene binding between CN females to PD females, PD SUB was higher than CN SUB by >250% (PD ♀ SUB/ CN ♀ SUB= 3.88) and in the case of HP, PD HP was higher than CN HP by >250% (PD ♀ HP/ CN ♀ HP= 3.81) (Figure 5B). When comparing [^18^F]nifene binding in all the subjects (males and females), PD SUB was higher than CN SUB by >300% (PD SUB/ CN SUB= 4.23) and in the case of HP, PD HP was higher than CN HP by >250% (PD HP/ CN HP= 3.80) (Figure 5C). Thus, a significant increase in [^18^F]nifene binding was measured in both males and females in the HP and SUB brain regions. Comparing the average of individual ratios of [^18^F]nifene binding between SUB/HP in PD (1.99) versus CN (1.67), there was a 19% increase in PD, suggesting that the SUB in PD was affected more than the HP and maybe because of the higher SUB binding in the males (Figure 5D).

**Figure 5.**
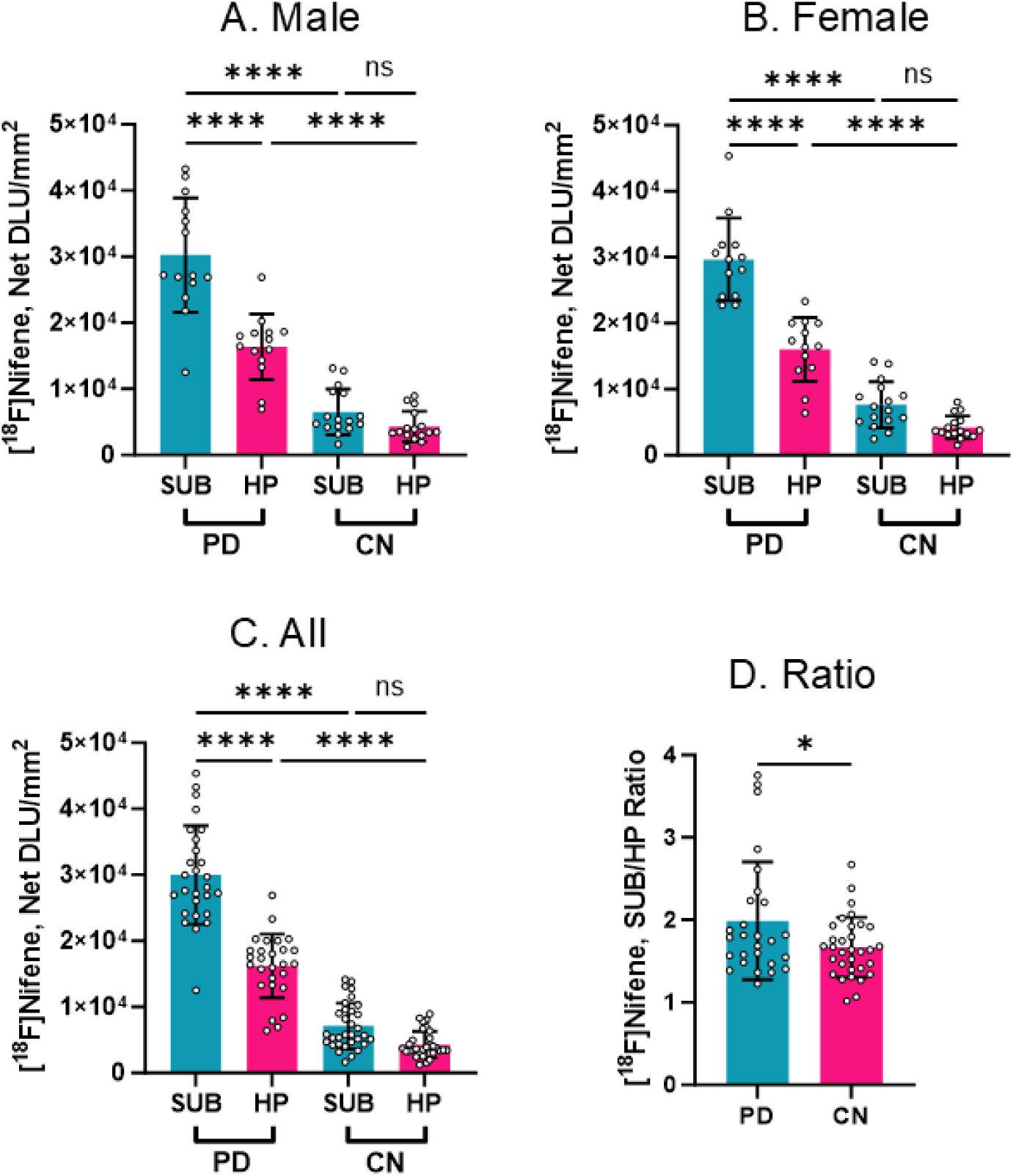
Comparisons between PD and CN [^18^F]nifene binding in SUB and HP. Unpaired two-tailed parametric t-tests determined statistical significance between each parameter (* *p* < 0.05, **** < 0.0001, ns = not significant). (A) [^18^F]Nifene binding in all male cases. (B) [^18^F]Nifene binding in all female cases. (C) [^18^F]Nifene in all male and female cases. (D) [^18^F]Nifene SUB/HP ratios of PD and CN.

### 3.4. Comparison of Grey Matter and White Matter

Along with the GM regions of SUB and HP, regions of WM were also quantified in all brain sections and compared to [^18^F]nifene binding in the GM regions which included SUB and HP (Figure 6). Comparisons of [^18^F]nifene binding between in GM between PD and CN cases were significant while WM comparison between PD and CN cases were not significant in both males and females (Figure 6A,B). In male comparisons, [^18^F]nifene binding to GM was greater than WM in CN by 37% (CN ♂ GM/WM= 1.37) and in PD by >250% (PD ♂ GM/WM= 3.63) (Figure 6A). When comparing [^18^F]nifene binding between CN males to PD males, PD GM/WM was significantly higher than CN GM/WM.

**Figure 6.**
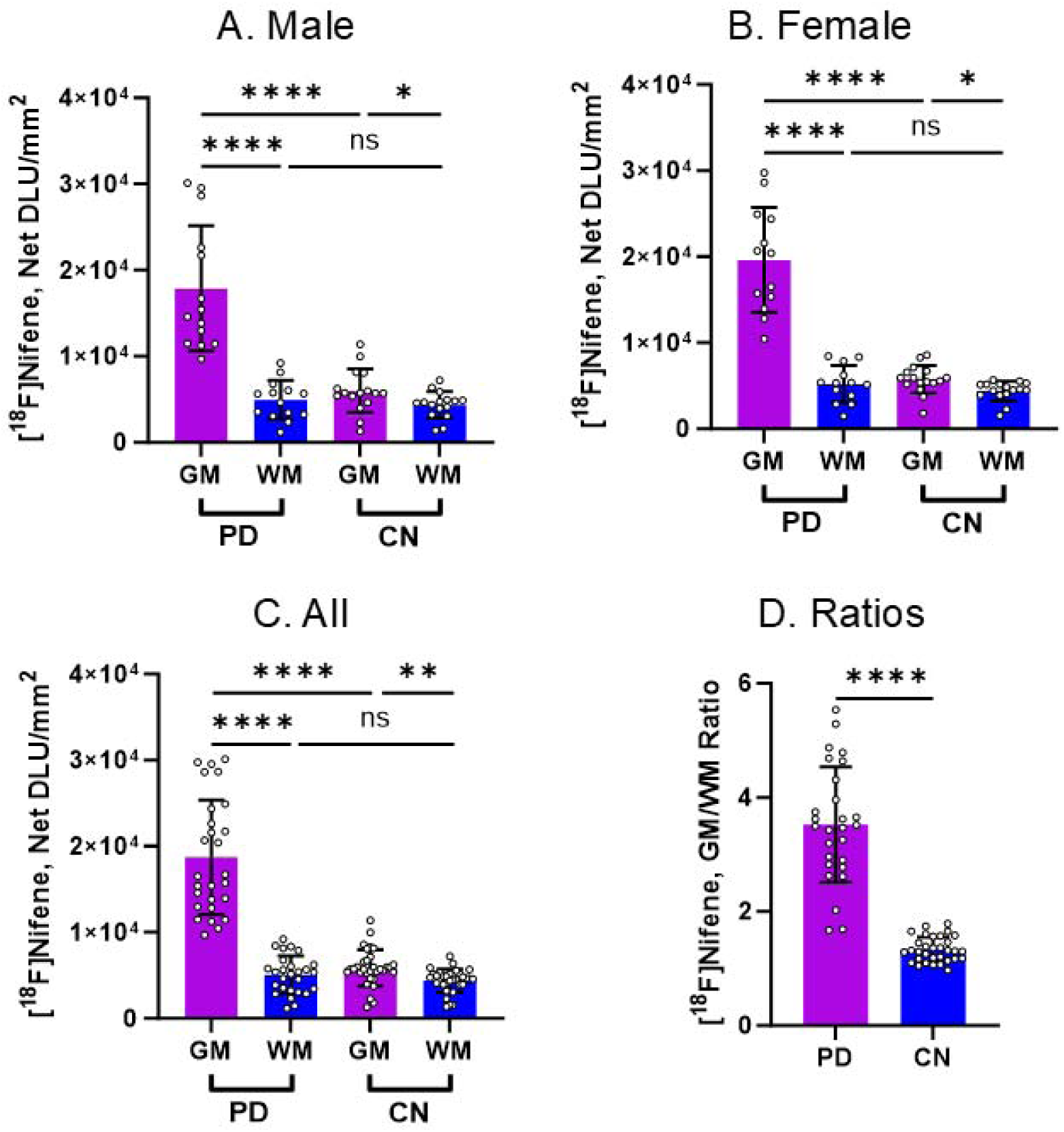
Comparisons between PD and CN [^18^F]nifene binding in GM and WM. Unpaired two-tailed parametric t-tests determined statistical significance between each parameter (* *p* < 0.05, ** p < 0.01, **** p < 0.0001, ns = not significant). (A) [^18^F]Nifene binding in all male cases. (B) [^18^F]Nifene binding in all female cases. (C) [^18^F]Nifene in all male and female cases. (D) [^18^F]Nifene GM/WM ratios of PD and CN.

In the case of females, comparisons of [^18^F]nifene binding to GM was greater than WM in CN by 30% (CN ♀ GM/WM= 1.30) and in PD by >250% (PD ♀ GM/WM= 3.76) (Figure 6B). Thus, similar to the males, female PD GM/WM was significantly greater than female CN GM/WM. The WM binding of [^18^F]nifene was very similar in the PD and CN cases (PD_WM_/CN_WM_=1.13 ♂;PD_WM_/CN_WM_=1.18♀) whereas GM binding was significantly greater in PD compared to CN (PD_GM_/CN_GM_=2.98 ♂; PD_GM_/CN_GM_=3.40♀). The GM/WM ratios across all PD subjects was 3.53 while the CN cases exhibited a ratio of 1.33, suggesting a >150% increase in the GM/WM ratio in PD (Figure 6D).

### 3.5. Aging Effects in PD and CN

Figure 7 shows the effect of age on [^18^F]nifene binding in CN cases across 3 decades in HP and SUB. Male cases exhibited an upward trend (increase in [^18^F]nifene binding in the GM) in both HP and SUB. Female cases on the other hand exhibited a small downward trend (decrease in [^18^F]nifene binding in the GM). When all the male and female cases were combined, there was little effect of aging in HP and SUB. A lack of aging effect was noted in human CN [^18^F]nifene PET study spanning over 6 decades (McVea et al., 2024).

**Figure 7.**
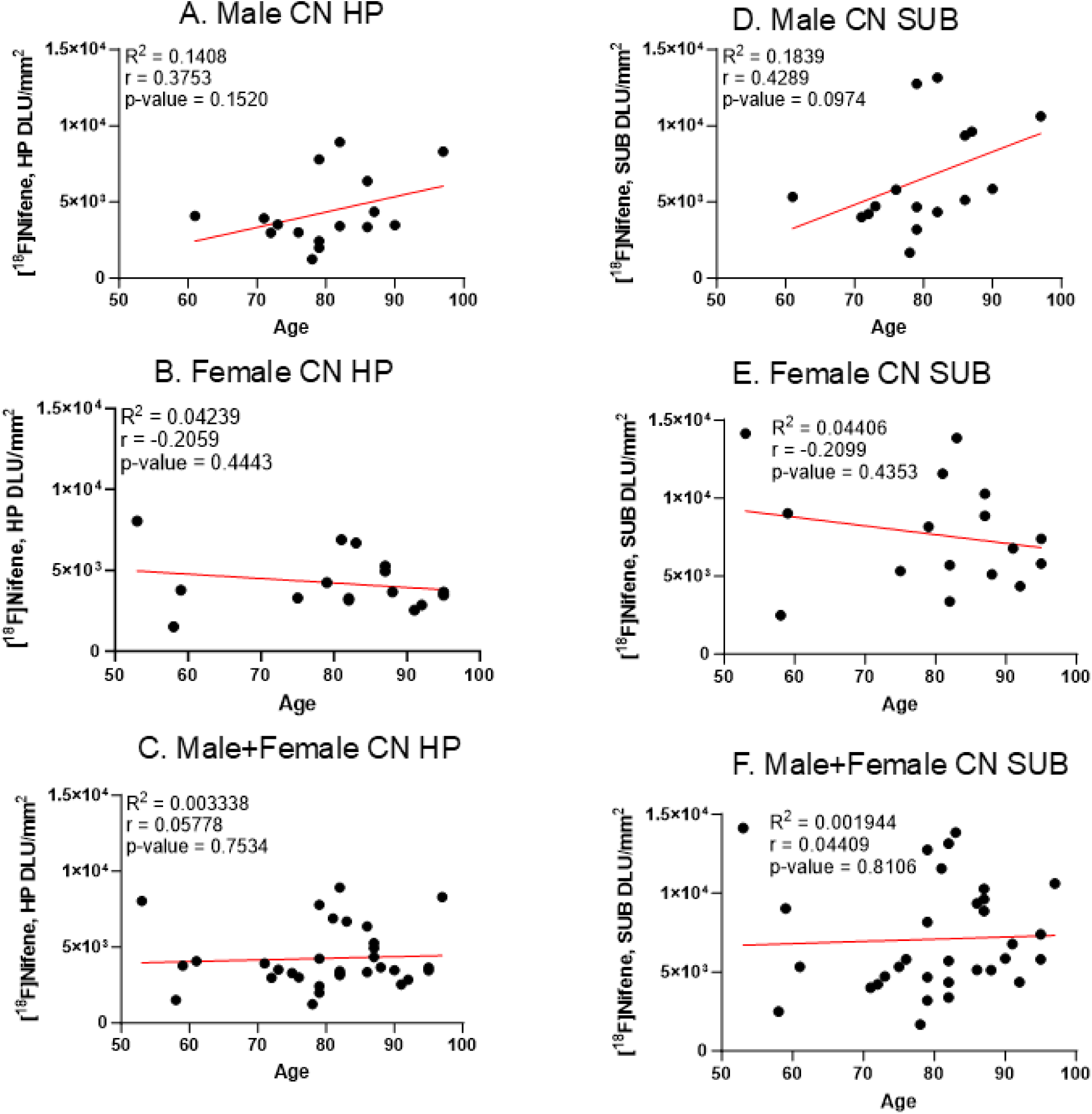
Aging effects of [^18^F]nifene binding in CN cases. (A). CN male HP, n=16 (Pearson’s r=0.3753; p value=0.1520); (B). CN female HP, n=16 (Pearson’s r= −0.2059; p value=0.4443); (C). CN male + female HP, n=30 (Pearson’s r=0.05778; p value=0.7534); (D). CN male SUB, n=16 (Pearson’s r=0.4289; p value=0.0974); (E). CN female SUB, n=16 (Pearson’s r= −0.2099; p value=0.4353); (F). CN male + female SUB, n=30 (Pearson’s r=0.04409; p value=0.8106).

In the case of PD, Figure 8 shows the effect of age on [^18^F]nifene binding to be different compared to the CN cases. Across 2 decades in HP and SUB. Male cases exhibited a downward trend (decrease in [^18^F]nifene binding in the GM) in both HP and SUB. The decrease was greater in the PD SUB compared to PD HP. Female PD cases on the other hand exhibited little change. When all the male and female cases were combined, there was a small downward trend in PD SUB [^18^F]nifene binding, while HP did not change significantly.

**Figure 8.**
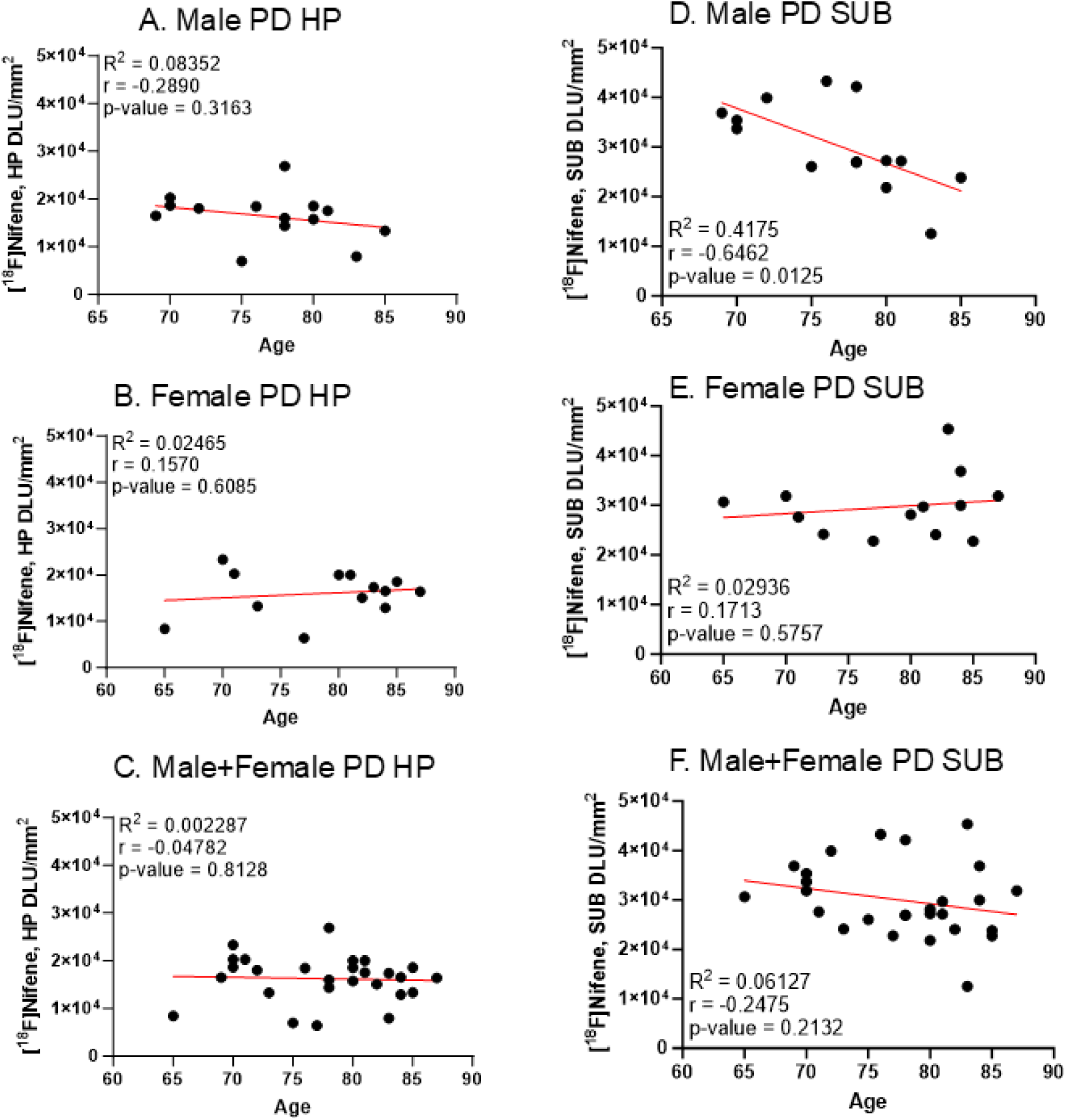
Aging effects of [^18^F]nifene binding in PD cases. (A). PD male HP, n=14 (Pearson’s r=0.2890; p value=0.3163); (B). PD female HP, n=13 (Pearson’s r= 0.1570; p value=0.6085); (C). PD male + female HP, n=27 (Pearson’s r=0.04782; p value=0.8128); (D). PD male SUB, n=14 (Pearson’s r= −0.6462; p value=0.0125); (E). PD female SUB, n=13 (Pearson’s r=0.1713; p value=0.5757); (F). PD male + female SUB, n=27 (Pearson’s r= −0.2475; p value=0.2132).

Thus, there appears to be differences of [^18^F]nifene binding between males and females in both CN and PD cases. In CN males, aging shows a slight increase in [^18^F]nifene binding in both HP and SUB, while in PD males, aging shows a slight decrease in [^18^F]nifene binding in both HP and SUB. The effect appears to be greater in the SUB compared to HP. In the females, differences were smaller, but the effect was reversed compared to the males. A downward trend was observed in CN females, while a very small increase was seen in PD females. Overall, there appears to be sex differences in [^18^F]nifene binding in both CN and PD cases.

### 3.6. Comparison of CN, PD and AD

Binding of [^18^F]nifene in the GM and WM regions was compared across PD, AD and CN cases (Figure 9A). Differences between GM and WM between the 3 groups were all significant. Binding of [^18^F]nifene was the lowest in AD and followed the order AD<CN<PD in both the GM and WM (Figure 9A,B). The AD GM was reduced by 32% compared to CN GM and AD WM was reduced by 40% compared to CN WM, whereas PD GM was higher than CN GM by >200% and PD WM was higher than CN WM by 26%. Figure 9C shows the GM/WM ratios of PD, AD and CN. The GM/WM ratios followed the order of CN<AD<PD with values of 1.33, 1.53 and 3.53 respectively. The higher WM [^18^F]nifene binding in the CN subjects changed the order of [^18^F]nifene binding when only GM was being considered.

**Figure 9.**
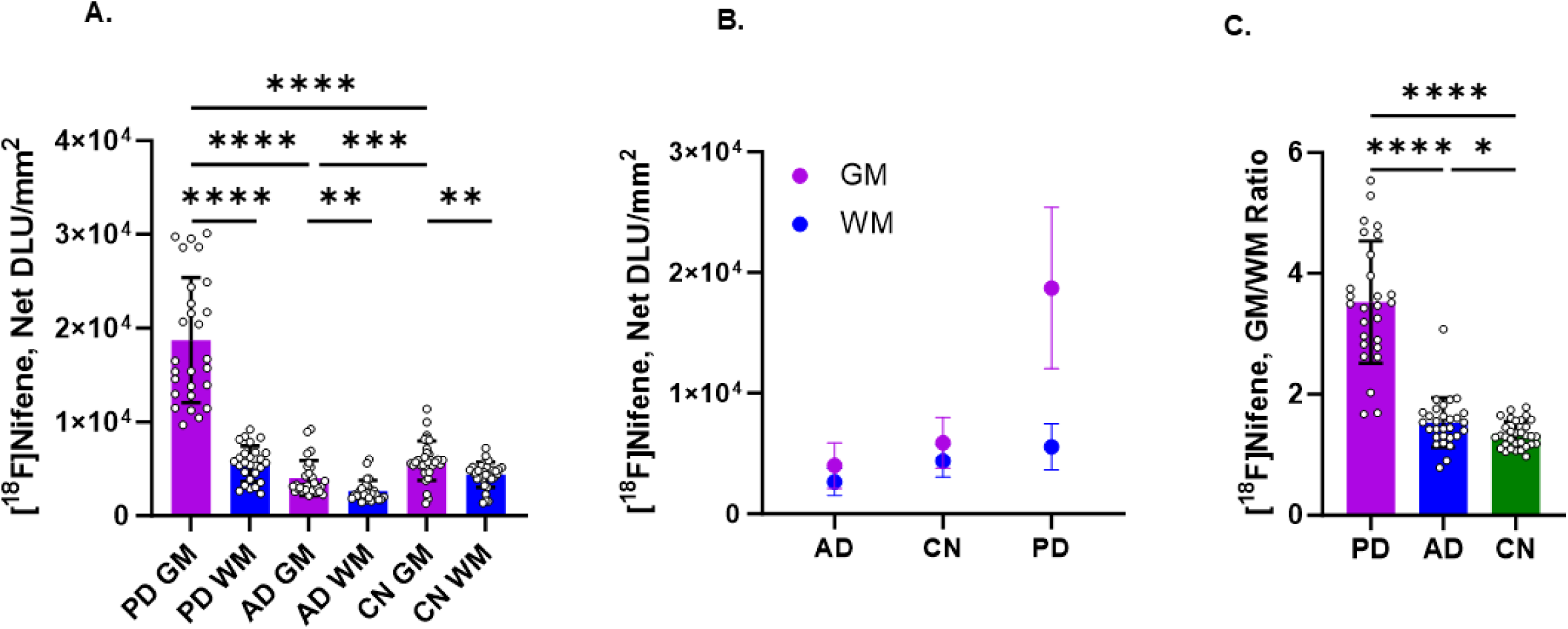
Comparisons of [^18^F]nifene binding between PD, AD, and CN. Unpaired two-tailed parametric t-tests determined statistical significance between each parameter (* *p* < 0.05, ** p < 0.01, **** p < 0.0001, ns = not significant). (A) [^18^F]nifene binding in GM and WM of PD, AD, and CN. (B) [^18^F]nifene binding to GM and WM superimposed per AD, CN and PD. (C) [^18^F]nifene GM/WM ratios of PD, AD, and CN.

## 4 Discussion

Classification of PD into 6 stages on the basis of α-synucleinopathy suggests the hippocampal region to be afflicted with α-synuclein aggregates in stage 4, causing cognitive deficits leading to dementia (Villar-Conde et al., 2021). As part of the subcortex, abnormalities in the hippocampus may play a role in SCAN which causes the diverse clinical manifestations of PD (Ren et al., 2026). Cholinergic neurotransmission in the hippocampus is maintained by different receptor subtypes which may be adversely affected in PD (Bohnen et al., 2022). In PD, cholinergic pathways are compromised as neurodegeneration progresses. To track how α4β2* nAChR affect PD pathology, [^18^F]nifene binding to α4β2* nAChR was evaluated in autoradiography. Using brain slices of only the HP-SUB in autoradiography allows clear visualization of the distinct subregions and differential [^18^F]nifene binding. Anti-ubiquitin and anti-α-synuclein immunostaining on adjacent slices shared distribution patterns with [^18^F]nifene binding in autoradiographic images. [^18^F]nifene binding was highly selective to α4β2* nAChR, more noticeably in the SUB than HP in both PD and CN. [^18^F]nifene binding in PD was significantly greater than CN in the SUB and HP regions, differing from [^18^F]nifene binding in AD which was less than CN.

Previous literature using other radiotracers for α4β2* nAChR had mixed results in greater or less binding in PD. A reduction in α4β2* nAChR was observed in PD striatum and substantia nigra using 2-[^18^F]FA-85380 PET (Meyer et al. 2009; Kas et al. 2009). Using 2-[^18^F]FA-85380 an increase in binding to α4β2* nAChR has been reported in HP (Meyer et al., 2005, 2009). Availability of α4β2* nAChR increased in the putamen, insular cortex, and orbitofronto-temporal cortices using 5-[^123^I]-A-85380 SPECT in cognitively intact early PD (Isaias et al. 2014). This was contrary to the decreases reported in several brain regions in non-demented PD subjects using 5-[^123^I]-A-85380 SPECT studies (<u>Fujita et al. 2005</u>; Oishi et al. 2007). [^18^F]XTRA binding was significantly higher in the PD occipital cortex, precuneus and mesial temporal cortex but not significant in HP (Mills et al. 2025). All these radiotracers were verified to bind to α4β2* nAChR but may differ in specificity and sensitivity that also depend on the type of subjects used. Additionally, cellular localization of α4β2* nAChR and the ability of the radiotracers to bind to different populations may play a role in some of the differences observed using different radiotracers (Zhang et al., 2023). Despite similar cholinergic denervation, vesicular acetylcholine transporter was upregulated exclusively in the HP among PD patients without cognitive impairment while PD with MCI was not (Legault-Denis et al. 2021). This hippocampal cholinergic upregulation suggests a compensatory mechanism that is unique to PD without cognitive impairment. Quantitative radioligand analysis in presymptomatic and early symptomatic mouse models revealed that nAChRs containing α4 and α7 participated in compensatory mechanisms that improved dopaminergic transmission (Kryukova et al. 2017; Albin et al. 2022). Brain regions may differ in degrees of cholinergic denervation that contribute to cognitive deficits. Accounting for the various affinities and sensitivities in PET tracers within specified subject characteristics allows for ensured accuracy in monitoring disease progression and therapeutic recovery.

Regardless of neurological condition, [^18^F]nifene binding was preferentially to α4β2* nAChR in the SUB. The mRNA of subunits α4 and β2 are strongly expressed the presubiculum, parasubiculum, subiculum, and dentate gyrus while moderately expressed in CA1-CA3 (Dineley-Miller & Patrick 1992; Drago et al. 2003). The distinction of subunit mRNA distribution patterns may explain preference for [^18^F]nifene binding to α4β2* nAChR within the SUB region. In the transgenic Hualpha-Syn(A53T) PD mouse model of α-synucleinopathy (Mondal et al., 2021), the SUB was among the brain regions with the highest amount of [^18^F]nifene along with the thalamus (Campoy et al. 2021). Similarly to the mouse model, the PD brain slices in this study contained an appreciable amount of α-synuclein.

Male-female differences in this study were significant. In the CN males, there was an upward trend of [^18^F]nifene binding with aging. In the case of PD males, this trend was lost and appeared to be on a downward trend with aging. It should be noted that in our [^18^F]nifene PET imaging study, there was no significant difference between the young and old CN subjects (McVea et al., 2024). Corpus callosum (CC) was used as a reference region in the [^18^F]nifene PET study. Previously CC has been reported to bind [^18^F]flubatine (Bhatt et al., 2018), however the degree to which [^18^F]nifene may bind to any available α4β2* nAChRs in the CC vis-à-vis aging is unclear (McVea et al., 2024). The females were distinctly different from males, both CN and PD were somewhat similar with little effect of aging. In the males, the SUB appeared to have greater increase in PD compared to females, whereas HP changes were similar for both males and females. Differences in non-motor cognitive impairment have been reported between males and females which may be related to neurochemical differences (Cerri et al., 2019; Cattaneo and Pagonabarraga, 2025).

Significant differences in [^18^F]nifene binding to α4β2* nAChRs in AD and PD were observed. Lower [^18^F]nifene binding in AD hippocampus-subiculum in postmortem brain slices was found in our recent study (Karim et al. 2025a) consistent with previous autoradiographic studies in AD using [^3^H]nicotine (Perry 1995). Binding of [^18^F]nifene was the lowest in AD and highest in PD and followed the order AD<CN<PD in both the GM and WM (Figure 9A,B). Presence of neuropathology in the AD HP-SUB (Aβ plaques, NFT and other inflammatory biomarkers) appear to have distinctively detrimental effects causing a decrease in [^18^F]nifene binding to α4β2* nAChRs. Neuronal loss in AD may be one of the reasons for this decrease in binding. Whereas in PD, accumulation of Lewy bodies and α-synuclein aggregates have the opposite effect resulting in increased [^18^F]nifene binding to α4β2* nAChRs. It is unclear if Lewy body may be the cause of this “accumulation” of α4β2* nAChRs in the cell body and/or if a-synuclein aggregate transport mechanisms from the presynapse may be causing an upregulation of α4β2* nAChRs resulting in increased [^18^F]nifene binding in PD. It is worth noting that the α7 nAChR binding measures using [^125^I]α-bungarotoxin in the HP-SUB regions of the same subjects were not significantly affected in both AD and PD (Ngo et al. 2025; Karim et al.,2025b).

Hippocampus in healthy subject PET studies of [^18^F]nifene showed a distribution volume ratio (DVR) of 1.44 using the corpus callosum WM as the reference region (McVea et al., 2024). This [^18^F]nifene DVR value of HP is in close agreement with the GM/WM ratio of 1.33 in control cases in the present study. The GM/WM ratio in the HP-SUB of PD cases in this study was 3.53 suggesting a significant increase in [^18^F]nifene α4β2* nAChRs. Recent studies using [^18^F]XTRA show increased binding in HP (Mills et al. 2025).

This study may be limited by possible variability between adjacent brain slices in IHC and autoradiography. Although the entirety of the HP-SUB region may not have been adequately visualized within brain slices of each case, the separation between SUB and HP was discernible. Quality of the autoradiographs minimally hindered analysis of [^18^F]nifene binding when drawing regions of interest and acquiring DLU/mm^2^ values. Regardless of the limitations, the results of this study support the use of [^18^F]nifene to measure α4β2* nAChR in PD. As a complimentary tool in diagnostics, [^18^F]nifene can assess the integrity of cholinergic function. This assessment can also encompass recovery of cholinergic function after therapeutically targeting α4β2 nAChR to ameliorate cognitive symptoms and pathology in PD (Quik & Wonnacott 2011; Liu et al. 2013). Previous validation of [^18^F]nifene in human PET and the results from this study encourage a seamless translation to future human PD PET studies.

## CONCLUSION

Overall, [^18^F]nifene exhibits binding to α4β2* nAChRs in human postmortem HP-SUB. A significant increase in the binding of [^18^F]nifene in PD suggests that this PET imaging agent can be used to reveal changes of α4β2* nAChRs as a diagnostic biomarker of PD. Further studies are needed to understand the influence of this elevated α4β2* nAChRs on SCAN in PD and potential for therapeutics development in PD (Goldman 2025).

## ACKNOWLEDGEMENTS

We are grateful to the Banner Sun Health Research Institute Brain and Body Donation Program of Sun City, Arizona for the provision of brain tissue. The Brain and Body Donation Program is supported by the National Institute of Neurological Disorders and Stroke (U24 NS072026, National Brain and Tissue Resource for Parkinson’s disease and related disorders), the National Institute of Aging (P30 AG19610 and P30AG072980, Arizona Alzheimer’s Disease Center), the Arizona Department of Health Services (contract 05700, Arizona Alzheimer’s Research Center), the Arizona Biomedical Research Commission (contracts 4001, 0011, 05-901 and 1001 to the Arizona Parkinson’s Disease Consortium) and the Michael J. Fox Foundation for Parkinson’s Research. We thank Jeffrey Kim, Pathology and Laboratory Medicine, University of California-Irvine for immunostaining of brain sections.

## Author Contributions

All authors had full access to all the data in the study and take responsibility for the integrity of the data and the accuracy of the data analysis. Conceptualization, J.M.; methodology, F.K., A.N., C.L., J.M.; G.E.S., T.G.B.; software, F.K., A.N.; validation and analysis, F.K., A.N.,C.L.; investigation, J.M..; resources, G.E.S, T.G.B.; writing—original draft preparation, F.K., J.M.; writing—review and editing, FK, JM, G.E.S, T.G.B.; supervision, J.M..; project administration, J.M..; funding acquisition, J.M.

## Data Sharing

The data that support the findings of this study are available from the corresponding author upon reasonable request.

